# Social complexity affects cognitive abilities but not brain structure in a Poecilid fish

**DOI:** 10.1101/2023.08.20.554009

**Authors:** Zegni Triki, Tunhe Zhou, Elli Argyriou, Edson Sousa de Novais, Oriane Servant, Niclas Kolm

## Abstract

Complex cognitive performance is suggested to be the out-turn of complex social life, allowing individuals to achieve higher fitness through sophisticated “Machiavellian” strategies. Although there is ample support for this concept, especially when comparing species, most of the evidence is correlative. Here we provide an experimental investigation of how group size and composition may affect brain and cognitive development in the guppy (*Poecilia reticulata*). For six months, we reared sexually mature female guppies in one of three different social treatments: (i) three female guppies; (ii) three female guppies mixed with three female splash tetras (*Copella arnoldi*), a species that co-occurs with the guppy in the wild; and (iii) six female guppies. We then tested the guppies’ performance in inhibitory control, associative learning and reversal learning tasks to evaluate their self-control, operant conditioning and cognitive flexibility capabilities. Afterwards, we estimated their brain size and the size of major brain regions using X-ray imaging technology. We found that individuals in larger groups of six individuals, in both same and mixed species treatments, outperformed individuals from the smaller groups of three guppies in reversal learning, with no apparent differences in the inhibitory control and associative learning tasks. This is rare evidence of how living in larger social groups improves cognitive flexibility, supporting that social pressures play an important role in shaping individual cognitive development. Interestingly, social manipulation had no apparent effect on brain morphology, but relatively larger telencephalons were associated with better individual performance in reversal learning. This suggests alternative mechanisms beyond brain region size enabled greater cognitive flexibility in individuals from larger groups.

## Introduction

Animals perform to remarkably high levels in various cognitive tasks, from simple associative learning to complex and sophisticated cognitive strategies. Nevertheless, there is tremendous variation in performance. Large-scale phylogenetic comparisons have revealed patterns and generated hypotheses why such variation exists; they show that species may evolve to adjust their brain morphology and, subsequently, their cognitive needs to their ecological conditions (Shultz and Dunbar 2006; van Schaik and Burkart 2011). Over decades, researchers have accumulated considerable knowledge about the various ecological factors (social and environmental) that may select for more sophisticated cognition and complex brains through genetic adaptation (evolutionary timescale). The social factors formed the basis for the social brain hypothesis (Dunbar 1998). This hypothesis states that brains (or brain regions) are enlarged due to selective social pressures linked to group size (Dunbar 1992, 1993; Barton 1996; Kudo and Dunbar 2001; Beauchamp and Fernández-Juricic 2004; Shultz and Dunbar 2006; Street et al. 2017), mating systems (Pawłowski et al. 1998; Iwaniuk 2001; Barton 2006), or social bonds (Emery et al. 2007; Dunbar and Shultz 2007). Although this is an evolutionary hypothesis that stems from between-species comparisons, it is possible also to use the logic of social pressures impacting brain morphology (Kotrschal et al. 2012; Fischer et al. 2015; Triki et al. 2019) and cognitive performance (Brown and Braithwaite 2005; Ashton et al. 2018; Triki et al. 2020) to explain variation in individuals’ strategies. This is supposed to help individuals to cope with their daily social challenges within lifetime, yielding an ontogenetic version of this hypothesis. Indeed, natural selection acting on existing genetic variation is not the only path by which individuals of a species adapt to their environment. Phenotypic plasticity, the interaction between an individual’s genome and its environment, is an additional path by which fine-tuned adaptations to local conditions are achieved (Kawecki and Ebert 2004; Sotka 2005).

Fishes emerge as a promising study clade for the ontogenetic approach as they possess more plastic brains than birds and mammals (Kotrschal et al. 1998), and possess relatively complex cognitive abilities (Triki et al. 2021; Bshary and Triki 2022). For example, wild cleaner fish living in high population densities grow larger forebrains than individuals living in low population densities (Triki et al. 2019), while lab-reared populations of sticklebacks housed in groups develop larger optic tectum compared to those housed individually (Gonda et al. 2009), and African cichlids experiencing early life rearing conditions in larger groups develop bigger hypothalamus and cerebellum and a different behavioural repertoire compared to those in smaller groups (Fischer et al. 2015). Additionally, adult female and male guppies living in same or mixed-sex pairs exhibit rapid sex-specific changes in brain morphology (Kotrschal et al. 2012). This indicates that fish can adapt to their environment quickly, making them perfect for testing the social brain hypothesis within an ontogenetic timeframe. How social pressures impact fish brain structure and subsequent cognitive abilities remains uncertain. Therefore, an integrative approach simultaneously investigating potential plastic changes in the brain and cognitive performance to adjust to social pressures in an experimental framework is pivotal to understanding cognitive development within the social brain hypothesis.

Here, we used the guppy (*Poecilia reticulata*) as a study system. In the wild, guppies live in shoals varying in size from only two fish to up to 50 per shoal, with frequent fission-fusion events allowing them to form complex and well-structured social networks (Croft et al. 2003, 2004). Also, guppies often coexist with, and may compete with, other fish species over resources (Anaya-Rojas et al. 2021). Currently, we need an understanding of how social challenges arising from competitive co-existence can impact animals’ cognitive and brain development. Therefore, in our experiment, we reared sexually mature female guppies in the same or mixed-species groups that varied in size for six months. Overall, we had three different social treatments, groups of three guppies living together, three guppies living with three other female fish of a different species, the splash tetra (*Copella arnoldi*) – a species that coexist with guppies in nature (Phillip 1998) – and six guppies living together. This allowed us to simultaneously test the effect of group size and same-species vs mixed-species compositions. We tested the guppies’ performance in three cognitive tasks: inhibitory control, associative learning and reversal learning. We estimated guppies’ inhibitory control abilities using a food reward placed inside a transparent cylinder. We recorded whether fish would delay their gratification and swim around and inside the cylinder without touching it – an indicator of inhibitory control ability – or whether they would bump into the cylinder in an attempt to retrieve the food immediately, which indicates a lack of inhibitory control. The cylinder test has been widely used to evaluate animal self-control capabilities (Kabadayi et al. 2018), from primates, and birds (MacLean et al. 2014), to fish (Lucon-Xiccato et al. 2017; Triki et al. 2023). For the associative learning task, we used a two-colour discrimination paradigm, where we evaluated guppies’ abilities in associating a colour cue (yellow or red) with a food reward and then reversed the colour-reward contingency (reversal learning) and estimated their performance in unlearning the previous rule and updating it with the current colour-reward association, an indicator of cognitive flexibility (Uddin 2021). Furthermore, we recorded fish behaviour to gain insight into how the social treatment may have impacted their social interactions. We focused on two traits: aggression and aggregation. This helped us determine if living in larger groups would lead to escalated conflicts and thus higher aggression rates. We also quantified aggregation to understand how the fish used the space in the different treatments. Afterwards, we acquired brain morphological measurements using state-of-the-art X-ray imaging technology (see Methods) to generate fine-tuned and high-quality volume data of fish major brain regions (White and Brown 2015): telencephalon, diencephalon, mesencephalon, cerebellum, and brain stem (Fig. 1).

**Figure 1.**
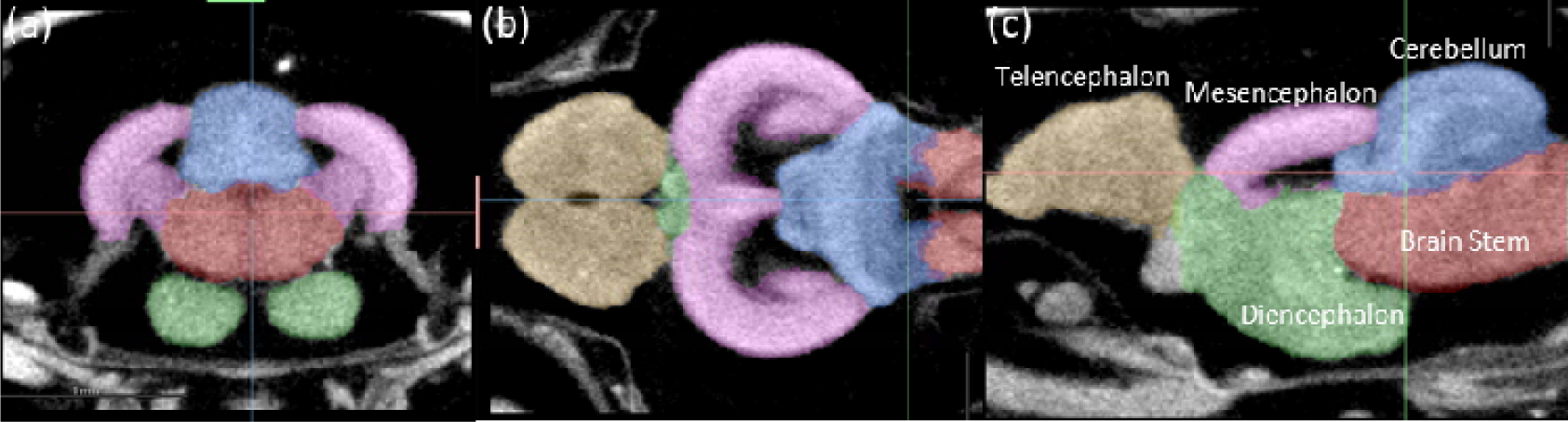
Segmented fish brain X-ray images. (a)Transversal, (b) coronal, and (c) sagittal planes of a brain scan. The major five brain parts were segmented and displayed in different colors: yellow – telencephalon, purple – mesencephalon, green – diencephalon, blue – cerebellum and red – brain stem.

If the social brain hypothesis applies to an ontogenetic timescale of individual development, our experiment should reveal a positive correlation between group size and cognitive performance in tests related to inhibitory control and reversal learning, potentially mediated by rapid changes in brain morphology, i.e., larger brains and region(s). This is based on our understanding of the social brain hypothesis at the evolutionary timescale, where species living in larger groups tend to have larger brains (Dunbar and Shultz 2007) and excel in complex cognitive tasks (Bond et al. 2003; Reader et al. 2011; Street et al. 2017). Furthermore, we expected this socially-mediated brain tissue expansion to yield better performance in our cylinder and reversal learning tasks. These predictions are based on comparative research indicating that species with larger brains (and brain regions) exhibit greater abilities in self-control and cognitive flexibility (Deaner et al. 2007; MacLean et al. 2014).

## Results

We raised 144 naïve female guppies in one of three different environments: a group of three guppies, a group of three guppies with three other fish of a different species (splash tetra), and a group of six guppies, resulting in 12 replicates per treatment. After six months, we sampled a subset of these guppies for X-ray brain imaging to estimate the overall size of their brains as well as five major brain regions. We tested the remaining guppies in three cognitive tasks, inhibitory control, associative learning and reversal learning tasks (Supplementary Videos). Upon finishing the tests, we sampled these fish for X-ray imaging and brain morphology analyses (see Methods). These fish were the core of our study to experimentally test the effect of social complexity on brain morphology and subsequent cognitive performance.

### Social treatment effect on cognitive performance

Among the three cognitive tests we ran, reversal learning emerged as the only one being affected by the social treatment (Generalized Linear Mixed Model using Template Model Builder (glmmTMB): N = 80, *X^2^* = 12.852, *p* = 0.002, explained variance: marginal-R^2^ = 0.35, conditional-R^2^ = 0.46). Fish from the treatment of six guppies and the treatment of three guppies with three splash tetras outperformed fish from the treatment of three guppies (posthoc test emmeans: six guppies vs three guppies, estimate = 1.971, *p* = 0.004; three guppies with three splash tetras vs three guppies, estimate = 1.309, *p* = 0.047), with no statistically significant differences between six guppies and three guppies with three splash tetras (estimate = 0.662, *p* = 0.584) (Fig.3). The other two tests, performance in the inhibitory control and associative learning tasks did not significantly differ across the three social treatments (glmmTMB: inhibitory control, N = 80, *X^2^* = 1.952, *p* = 0.376; associative learning, N = 80, *X^2^* = 1.611, *p* = 0.446) (Fig. 2).

**Figure 2.**
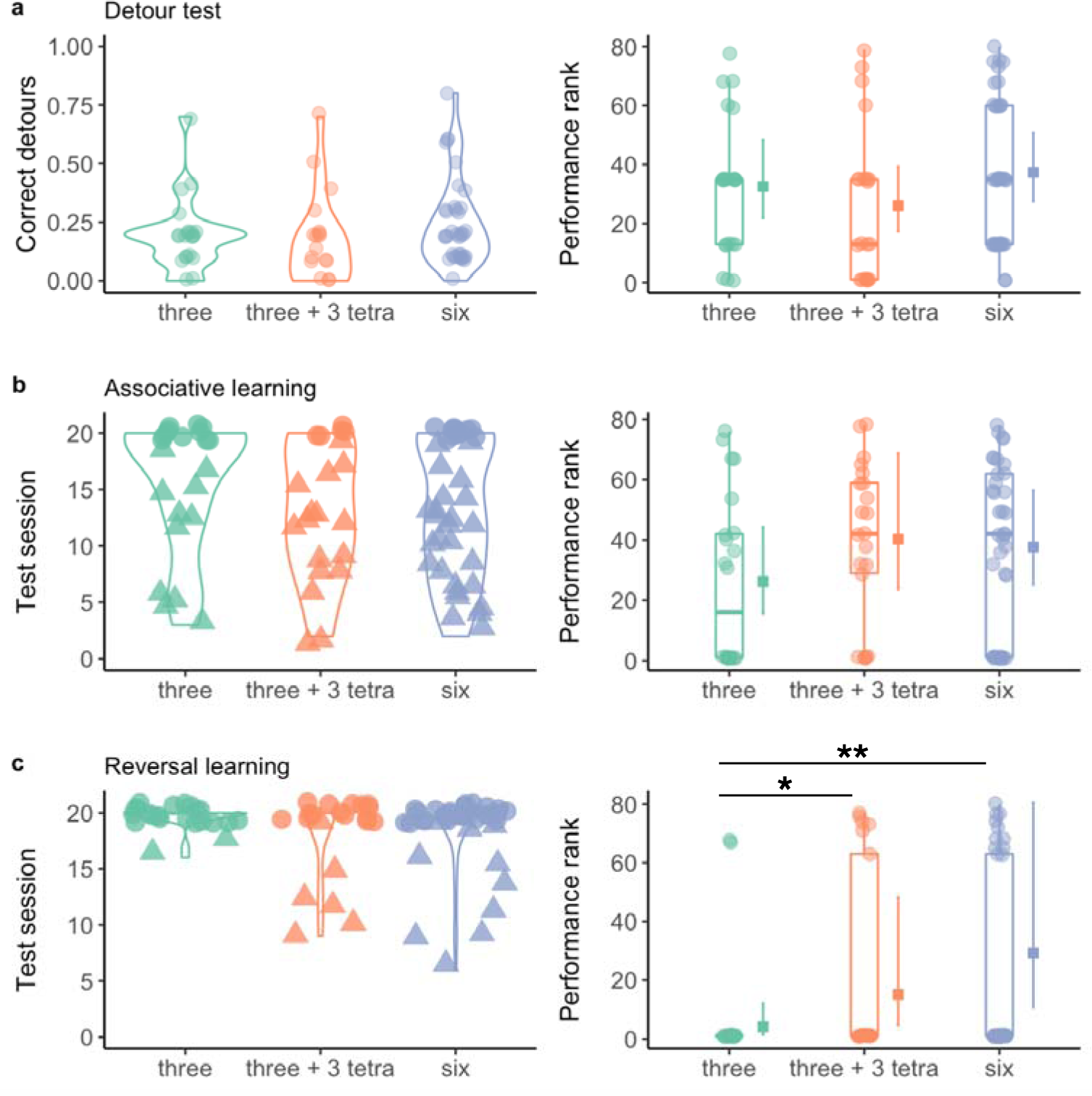
Cognitive performance of the guppies from the three social treatments. The panels on the left show the raw scores per cognitive task as violin plots with data points, while panels on the right show the estimate and 95% CI of model marginal effects, combined with boxplots of median and interquartile of performance rank (N = 80) for (a) Inhibitory control (b) associative learning and (c) reversal learning. The highest ranks refer to the highest performance. The three social treatments on the X-axes are social groups of three guppies, three guppies with three splash tetras, and six guppies. The reversal learning test shows an effect of social treatment on fish performance (\**p* < 0.05, \*\**p* < 0.01), but not the other two tests (*p* > 0.05).

**Figure 3.**
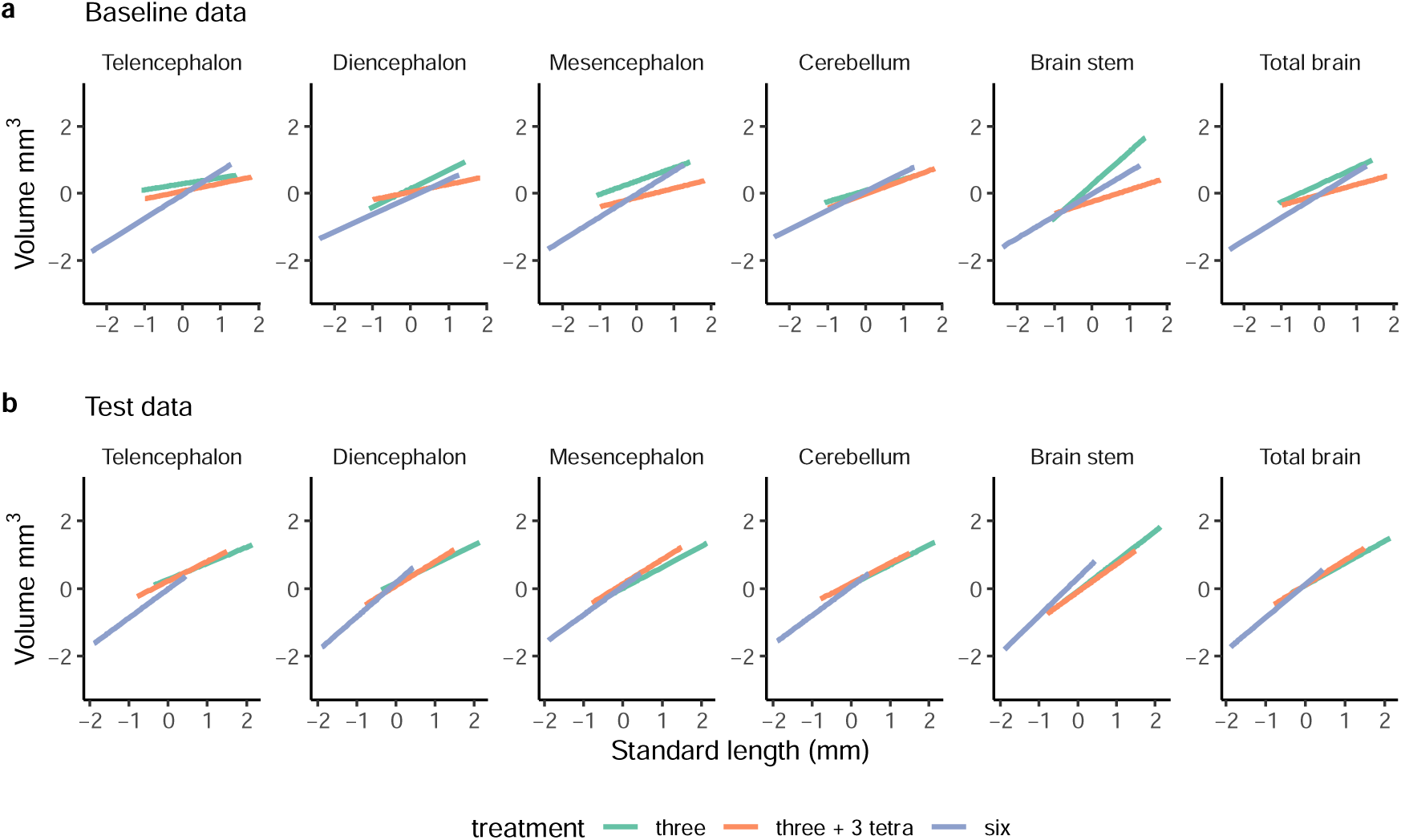
Brain morphology of the guppies from the three social treatments. Scatterplot and regression lines of log-normal transformed and standardized volume (mm^3^) of the brain measurement on the log-normal transformed and standardized body size (standard length in mm) as a function of social treatment, from (a) the baseline dataset of the 29 female guppies sampled before the cognitive tests and (b) test dataset (80 female guppies) after the cognitive tests. The three social treatments indicated by different colours are social groups of three guppies, three guppies with three splash tetras, and six guppies. There were no significant effects of social treatment on brain morphology (*p* > 0.05).

### Social treatment effect on qualitative behavioural traits, aggression and aggregation

The detailed statistical outcomes of the behavioural analyses are reported in Supplementary Table S1. Overall, we found that the social treatment, to some extent, affected the aggression rates towards guppy fish (LMM: *X^2^* = 6.198, *p* = 0.045, marginal-R^2^ = 0.18, conditional-R^2^ = 0.20, see Supplementary Material), apparently driven by the group of mixed species where splash tetras were the most aggressive.

### Social treatment effect on brain morphology

Guppies’ brain morphology did not change as a function of social treatment. Neither fish sampled before (N = 29) nor those sampled after (N = 80) the cognitive tests showed a significant (*p* > 0.05) change in overall brain size or the five regions quantified here (telencephalon, diencephalon, mesencephalon, cerebellum, and brain stem) (Fig. 3) (see detailed statistics in Table 1). All measurements were corrected for body size since body growth across treatments varied significantly. Guppies from the treatment of six guppies were significantly smaller than guppies in the treatment of three guppies and the treatment of three guppies with three splash tetras (linear mixed effects model (LMM): from baseline data: *X^2^* = 7.843, *p* = 0.019, R^2^ = 0.22; from test data: *X^2^* = 129.5, *p* < 0.001, marginal-R^2^ = 0.55; conditional-R^2^ = 0.66, see Supplementary Material).

**Table 1.** Summary table of the outcomes of brain morphology as a function of social treatment. Values in bold refer to statistically significant results with a p-value ≤ 0.05.

| Dataset | Explanatory variable | test | N | F-value | p-value |
| --- | --- | --- | --- | --- | --- |
| Total brain size |  |  |  |  |  |
| Baseline | Social treatment | LMM | 29 | 1.458 | 0.482 |
|  | Body size |  |  | **13.421** | **<0.001** |
| Test | Social treatment | LMM | 80 | 0.359 | 0.835 |
|  | Body size |  |  | **59.764** | **<0.001** |
| Telencephalon, diencephalon, mesencephalon, cerebellum, and brain stem sizes |  |  |  |  |  |
| Baseline | Social treatment | MANOVA | 29 | 0.362 | 0.955 |
|  | Body size |  |  | 2.542 | 0.065 |
|  | Batch identity |  |  | 1.920 | 0.140 |
|  | Social treatment x batch identity |  |  | 0.389 | 0.943 |
| Test | Social treatment | MANOVA | 80 | 0.908 | 0.527 |
|  | Body size |  |  | **14.598** | **<0.001** |
|  | Batch identity |  |  | **2.344** | **0.050** |
|  | Social treatment x batch identity |  |  | 0.499 | 0.887 |

### The link between cognitive performance and brain morphology as a function of social treatment

By looking at individual performance in every test (Fig. 4-6) and correlating it to brain morphology and social treatment, we found that relative telencephalon size was positively associated with improved performance in the reversal learning task (glmmTMB: N = 80, telencephalon size residuals, *X^2^*= 6.374, *p* = 0.011, marginal-R^2^ = 0.45, conditional-R^2^ = 0.55, Fig. 6), but independently of social treatment (glmmTMB: N = 80, telencephalon size residuals x social treatment, *X^2^* = 1.319, *p* = 0.517) (Fig. 4). For the other brain measurements as well as the other two cognitive tasks, i.e., inhibitory control and associative learning, we did not find any statistically significant relationships (Fig. 4, see detailed statistics in Table 2).

**Figure 4.**
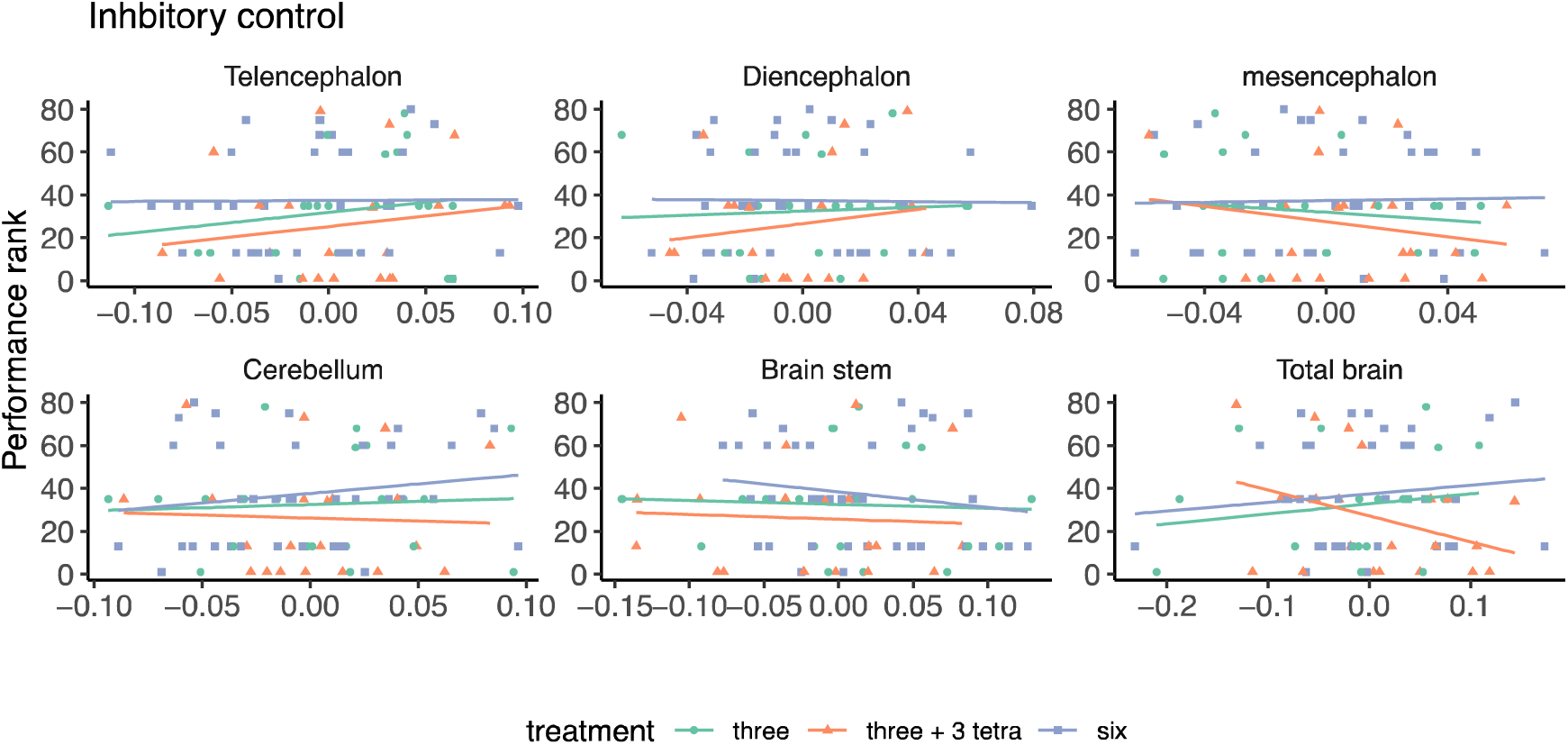
The relationship of inhibitory control performance and brain morphology of the guppies from the three social treatments. Scatterplot and regression lines of performance rank (where highest ranks refer to highest performance) on brain measurement residuals (x-axes) (N = 80). The three social treatments indicated by different colours are social groups of three guppies, three guppies with three splash tetras, and six guppies. No significant effect was detected (*p* > 0.05).

**Figure 5.**
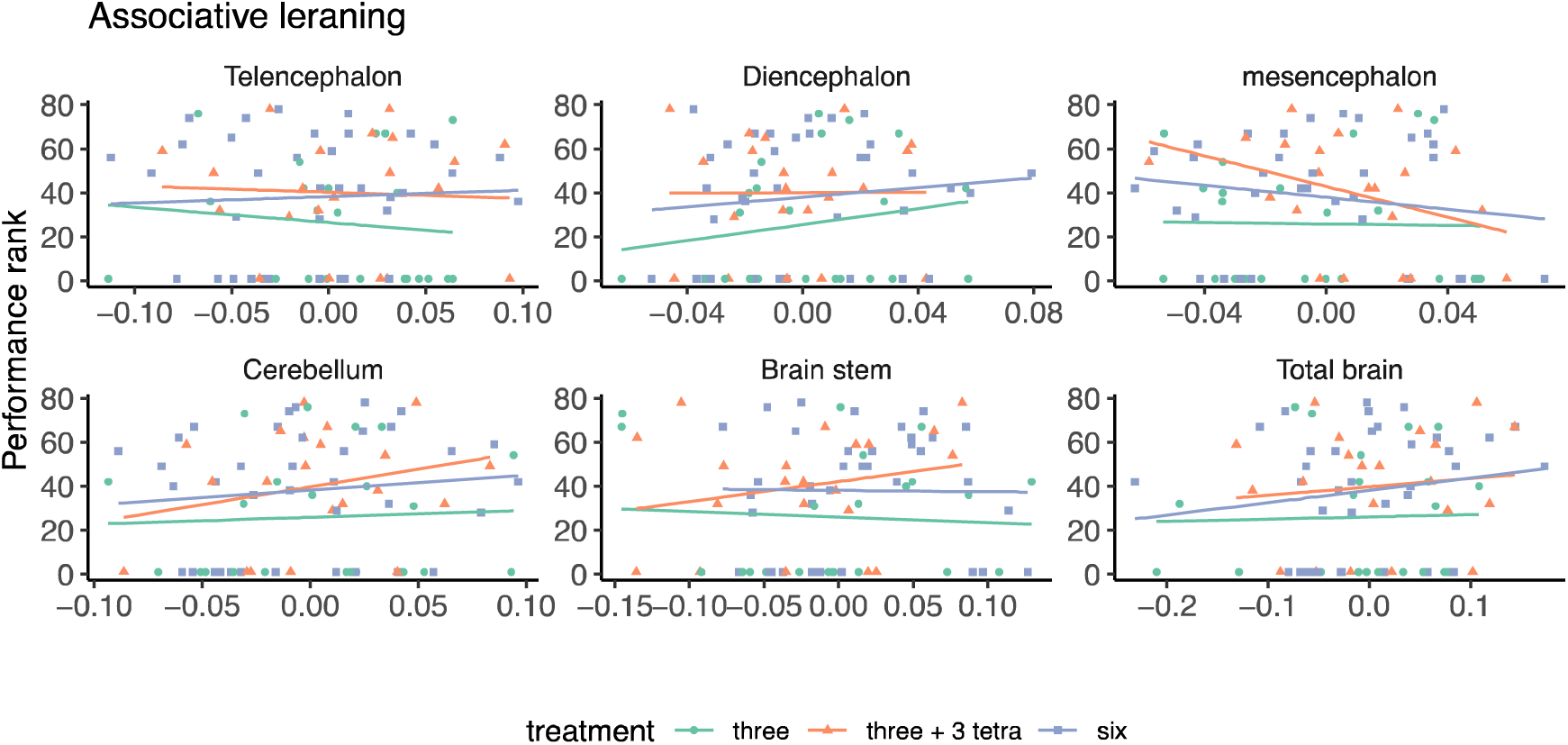
The relationship of associative learning performance and brain morphology of the guppies from the three social treatments. Scatterplot and regression lines of performance rank (where highest ranks refer to highest performance) on brain measurement residuals (x-axes) (N = 80). No significant effect was detected (*p* > 0.05).

**Figure 6.**
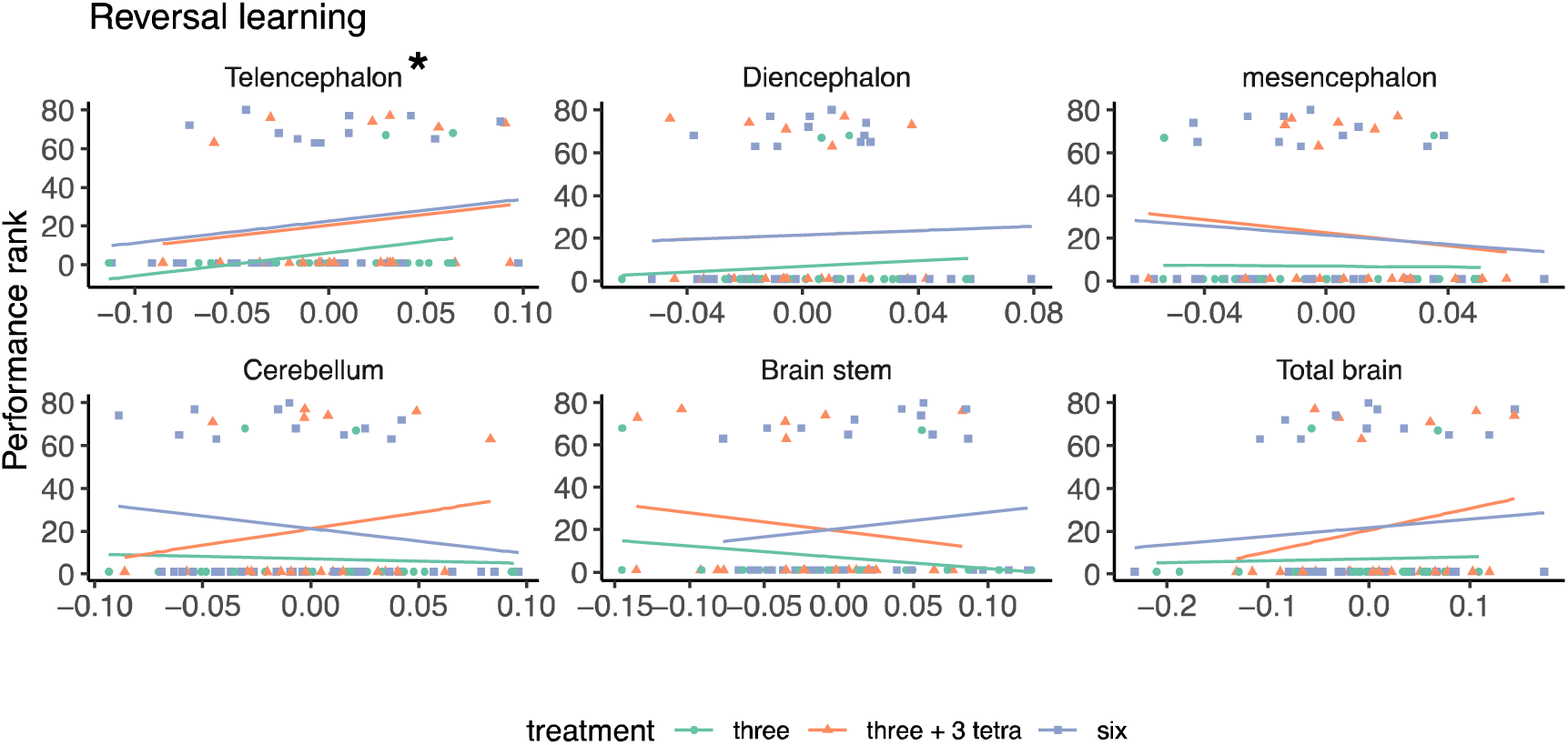
The relationship of reversal learning performance and brain morphology of the guppies from the three social treatments. Scatterplot and regression lines of performance rank (where highest ranks refer to highest performance) on brain measurement residuals (x-axes) (N = 80). Only the telencephalon relative size correlated significantly with performance (*p* < 0.05).

**Table 2.**
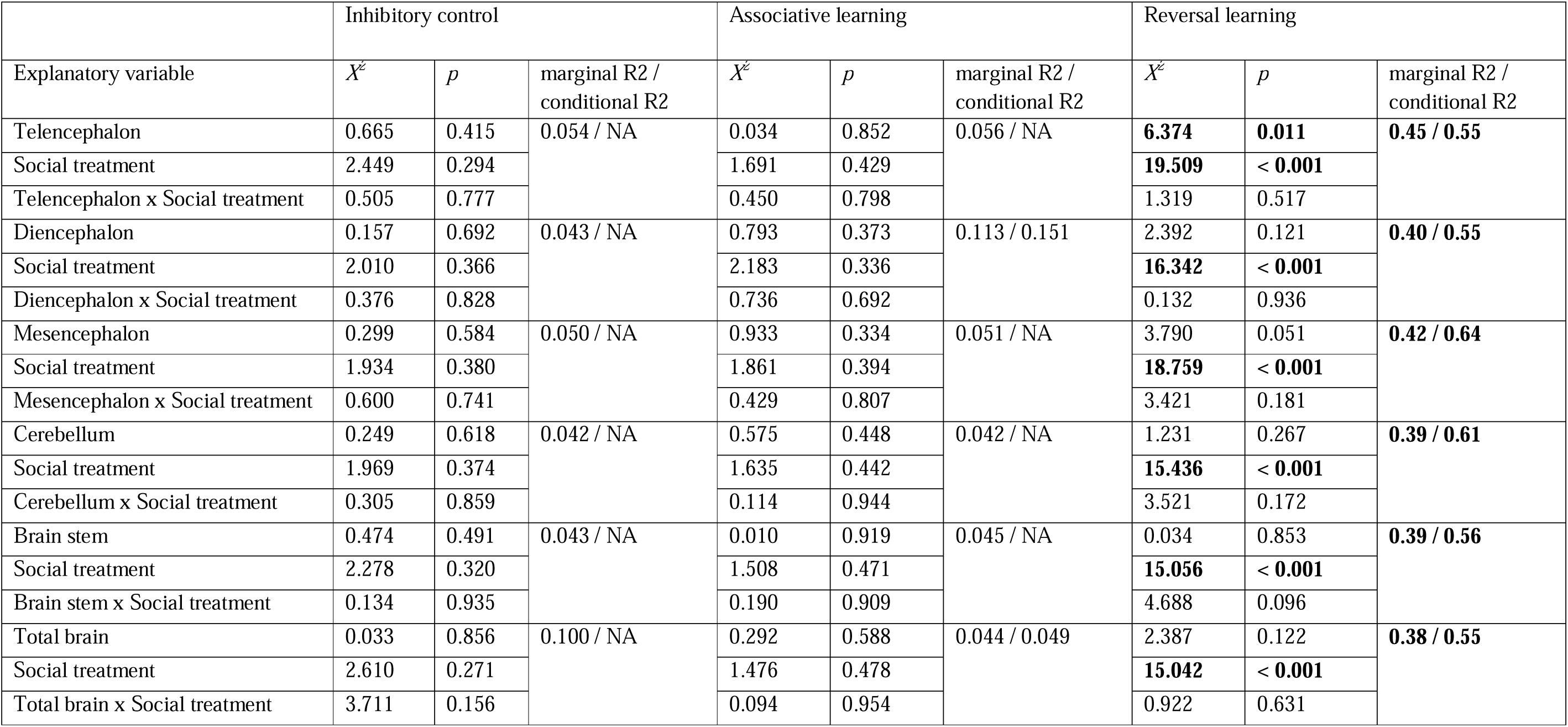
Summary table of the outcomes of cognitive performance as a function of brain morphology and social treatment. Values in bold refer to statistically significant results with a p-value ≤ 0.05. Sample size is N = 80 guppies.

## Discussion

Our study tested the social brain hypothesis within an ontogenetic timescale. We asked whether social group size and group composition affect brain morphology and cognitive performance across different cognitive domains in a guppy species. The key findings were: (i) living in large groups of six individuals, either of the same or mixed species, yielded improved performance in the reversal learning task than those in small groups of three individuals, (ii) social treatment did not affect associative learning performance and inhibitory control, (iii) social treatment did not affect brain morphology, and (iv) independently of social treatment, relative telencephalon size correlated positively with individual performance in the reversal learning task. We discuss each finding in turn in the following paragraphs.

Living in larger groups can create richer social environments and lead to the development of more sophisticated strategies to cope with daily challenges (Dunbar and Shultz 2007). This, in turn, can result in more advanced cognitive abilities. Our study found that guppies living in groups of six individuals exhibited better cognitive flexibility, as expected. This aligns with previous studies showing a positive correlation between social complexity and cognitive flexibility (Byrne and Whiten 1989; Bond et al. 2007; Ashton et al. 2018). In larger groups, individuals must be able to switch attention and adjust behaviours with changing demands more effectively than when cohabiting with fewer individuals. Interestingly, our study found that regardless of whether a guppy lived in larger groups of the same or a different species, their cognitive capacities developed similarly; a finding that is comparable to that by Fischer et al. (2021) on cichlids. It seems that the amount of social interactions that occur when living in larger groups is more important for cognitive flexibility than the exact nature of these interactions. Alternatively, the differences in social interactions between the two species included here might not be large enough to generate differences in the cognitive tasks we quantified. Indeed, our analysis of specific behaviours, specifically aggression and aggregation, did not provide any clear explanations for the differences observed in group performance during the reversal learning task. This result shows that to understand the influence of the social environment on cognitive development, we may have to use a more holistic ecological framework, including also social interactions across species. It would be highly interesting to extend this finding and also include social interactions with other species in comparative phylogenetic analyses of the link between group size and composition and cognitive evolution. Beyond the predator-prey interactions (Bijl and Kolm 2016), this has not been done yet. Moreover, an important applied outcome of this result is that social enrichment could be highly effective also when created through the inclusion of other species, for instance, in zoos and sanctuaries (Dorman and Bourne 2010).

Unsurprisingly, there were no differences in associative learning performance due to social treatment. The test assesses basic operant conditioning abilities, and simple cognitive processes are sufficient for forming associations (Savage 1980; Triki et al. 2023). The task is often excluded when researchers look into complex cognitive abilities, like general intelligence (Damerius et al. 2017; Aellen et al. 2022). In sum, social complexity would unlikely impact how an individual forms basic associations, such as between a colour cue and a food reward.

The ability to pause and override motor impulses in response to a specific stimulus is known as inhibitory control. When executed correctly, this results in adaptive, goal-oriented behaviours requiring complex cognitive processes (Diamond 2013). We used the detour task to test fish performance in this cognitive capacity, and the results showed that social treatment did not affect their inhibitory control abilities. This finding goes against our original predictions as previous studies have shown that living in complex social environments enhances inhibitory control abilities for individuals and species (Amici et al. 2008; Ashton et al. 2018; Johnson-Ulrich and Holekamp 2020). However, it is worth noting that other ecological factors, besides social complexity, may also have significant impacts (see review by Rosati (Rosati 2017)). Currently, we rely heavily on comparative and correlative research to study the relationship between social complexity and inhibitory control. Therefore, we need more experimental data at both the species and population levels to draw convincing conclusions about whether social complexity directly enhances inhibitory control capabilities.

Regarding the brain morphology analysis, there were no evident changes caused by the social treatment. Still, we noticed that the brain allometry was different, with fish from the treatment of six guppies having relatively steeper allometry slopes for overall brain size but also for most of the brain regions on body size, compared to the other two treatments (Fig. 2). It is clear that differences in body growth drove this. Although we fed all fish *ad libitum* and we expected that their body growth would be density-dependent (Lorenzen and Enberg 2002), there were differences across treatments, where guppies in the small group of three and the large group of mixed species were of similar body sizes, but they were substantially larger than those in the group of six guppies. It suggests that guppies were more successful foragers than splash tetras in the groups of three guppies and three splash tetras because they attained larger body sizes as if they were alone in the tank compared to those growing relatively smaller when they were competing against their conspecifics in the groups of six guppies. Despite these body growth differences across treatments, there were no evident changes in brain size or the size of the five major brain regions when corrected for body size. We can only speculate on why we did not find differences here, when other studies on fish have found substantial responses in relative brain region size to variation in the social environment. For instance, cleaner fish living at higher population densities possess larger forebrains (telencephalon and diencephalon) (Triki et al. 2019), while nine-spined sticklebacks reared in groups develop a larger optic tectum and a smaller olfactory bulb than those reared individually (Gonda et al. 2009) (see review by Gonda et al. (2011) for more examples and detailed discussion). It is possible that we did not see any changes across different social treatments because increasing the group size from three to six was not enough to create the necessary social effects that lead to brain size and region sizes change. Yet, it is also possible that the treatments generated effects on a different scale that could not be detected with our X-ray scan methods. For instance, it could be that changes in neural connectivity, neuronal activity or gene expression, while not essentially leading to volume changes, were responsible for the observed group performance differences (Weitekamp and Hofmann 2014; Herculano-Houzel 2017; Wallace and Hofmann 2021). Another possible reason for the lack of social treatment effect on brain morphology in our data is that we only manipulated group size, ignoring other key factors that exist in the wild and affect brain development, such as predation pressure, mating strategies, and feeding ecology (Brown and Braithwaite 2005; Kolm et al. 2009; Powell et al. 2017).

We found that relative telencephalon size explained, to some extent, fish performance in the reversal learning test with no apparent differences in this brain region volume across social treatments. Specifically, relative telencephalon size correlated positively with reversal learning performance within each social treatment (see Fig. 4c), but it did not explain performance differences across treatments. Individual-level improvement in reversal learning performance due to larger telencephalon has already been demonstrated in guppies by Triki et al. (2022b, 2023). Additionally, Triki et al. (2022a, 2023) found a positive correlation between the size of this brain part and inhibitory control abilities, which was not observed in the current study. One possible reason for this could be that Triki et al.’s studies (2022a, 2023) involved the use of guppies that were selectively bred on divergence in relative telencephalon size over several generations. Often, it is difficult to detect brain morphology effects on behaviour in wild-type strains of laboratory-held animals that are fed *ad libitum* and where predation selection pressures are removed (see discussion on this topic in (McNeil et al. 2021). Hence, while not general across all cognitive abilities assayed here, we find it interesting that the effect of relative telencephalon size is consistent both for wild-type guppies and for artificial selection line guppies targeted for telencephalon size.

In summary, our research suggests that social complexity affects cognitive flexibility but not inhibitory control or basic operant conditioning skills. This impact is likely driven by mechanisms beyond plastic changes in brain region sizes. More research is necessary, using experimental methods, to fully understand how social and environmental factors shape cognitive development.

## Materials and methods

### Study animals and experimental set-up

We conducted the study between December 2019 and January 2021 in the fish laboratory facilities at Stockholm University in Sweden. Our study animals were laboratory-bred guppies *Poecilia reticulata* descendants from an initial population of more than 500 fish caught from high predation risk sites in 1998 in Quare river, Trinidad. To create a new generation of naïve guppies, we set up 75 breeding pairs in separate 2 L tanks. We regularly isolated the fry and housed them in 2 L tanks with a maximum capacity of six per tank. We periodically removed the ones that developed into males by showing secondary sexual traits like colour patterns, housed them in separate tanks, and ensured that the remaining female numbers were adjusted to six per tank. On average, guppies reach sexual maturation within three months of age. We also used another fish species, the splash tetra, *Copella arnoldi*, an indigenous species to the native Trinidadian rivers where the guppy typically resides. We obtained the splash tetras from an aquarium fish supplier in Stockholm. Finally, we used 144 female guppies from our new generation and 36 female splash tetras (divided into two batches) to create three social treatments with different fish densities and compositions.

The first treatment had three female guppies in one tank, the second treatment had three female guppies with three female splash tetras in one tank, and the third treatment was six female guppies in one tank. We used only females to avoid mating and reproduction occurring and potential male-male aggression. Every treatment had 12 replicate tanks, but some replicates had a later development of apparent males, whose colour patterns had not yet developed when we established the treatments. This led to the elimination of one replicate in the treatment of “three guppies”, one in the “three guppies with three splash tetras”, and four in the “six guppies”. After that, the sample size was 33 fish (11 tank replicates) in “three guppies”, 33 fish (11 tank replicates) in “three guppies with three splash tetras”, and 48 fish (8 tank replicates) in “six guppies” treatments. All housing tanks were of 6 L capacity and contained identical enrichment of 2 cm of gravel, one plastic plant in the middle and an air filter (see Supplementary Material). We ensured an ambient temperature of ∼ 26 °C and a light:dark cycle of 12:12 h with an *ad libitum* feeding schedule alternating between fish flakes and live *Artemia* (brine shrimp) hatchlings six days per week.

After six months, we terminated the three social treatments and allocated our focal individuals, i.e., the female guppies, to a “baseline” and “test” set while transferring the splash tetras to a 200 L housing tank. The baseline set served to test for brain morphology changes right after the termination of the social treatment but before the cognitive tests. In contrast, we subjected the test set first to a battery of cognitive tasks and then measured their brains (see below). In the baseline set, there were ten fish from “three guppies”, ten from “three guppies with three splash tetras”, and nine from “six guppies” treatments. The remaining fish (23 from “three guppies”, 23 from “three guppies with three splash tetras”, and 39 from “six guppies” treatments, see Supplementary Material) were housed individually in experimental aquaria (L × W × H; 40 × 15 × 15 cm). To avoid potential observer bias, the test fish had running number labels (1, 2, 3, etc.) to conceal their social treatment identity throughout the experiment. Each experimental aquarium had an identical enrichment of 2 cm of gravel and an artificial plant and had continuously aerated water; it also had two adjacent guillotine doors, one see-through and one opaque, dividing the space into a housing compartment and a test compartment. The experimental room had an ambient temperature of ∼26°C with a light schedule of 12 h light and 12 h dark. We fed the guppies *ad libitum* with defrosted adult brine shrimps delivered in a 1 mL transparent plastic pipette six days per week. This facilitated acclimating the fish to receive food from plastic pipettes, later used to provide food as a positive reinforcement in the learning tests. During the weekdays, when we ran cognitive tasks, fish acquired food solely from test trials. The tests started after an acclimation period of five days, with no trials on the weekends. During the tests, the between-trial interval for every fish was about 60 minutes, and one test trial per fish took about 1 min. Furthermore, there was always at least one day break between every two cognitive tests. Unfortunately, five fish, out of 85, died during the experiment after jumping out of the experimental tanks during the night. That left 22 fish from the treatment “three guppies”, 21 from the “three guppies with three splash tetras”, and 37 from the “six guppies”.

Additionally, while running the three social treatments, we attempted to acquire behavioural observations to establish a basic understanding of the social interactions occurring in these treatments. Within our logistic capacities, we recorded fish behaviour using video recording in 17 tanks, six tanks from “three guppies”, six from “three guppies and three splash tetras”, and five from “six guppies” tanks from the second batch after 5 months of treatment. The Supplementary Material contains further information on how we performed these observations and the resulting statistics.

### Cognitive tests

#### 1- Inhibitory control task (detour task)

After the acclimation period, we trained fish to associate the colour green with a food reward. To do so, we placed a green disc in the test compartment and delivered a defrosted adult artemia placed right on top of the disc. We repeated this exposure twice a day for four consecutive days. On the following day, we introduced a transparent Plexiglas cylinder (5 cm in length and 4 cm L) open on either side in the test compartment. The cylinder contained a food reward placed on top of a green spot drawn inside the cylinder. This was a one-time acclimation opportunity for the fish to explore the transparent barrier. After that, fish received three trials on test Day1, three trials on Day2 and four trials on Day3. A trial started when the experimenter pulled up the opaque and transparent guillotine doors simultaneously and allowed the fish to detour the physical barrier, here as the cylinder walls, and swim inside the cylinder to retrieve the food reward. The experimenter recorded whether the fish touched the cylinder (failure) or not (success) before retrieving the food (see Supplementary Videos). Finally, we ranked the proportion of correct detours (the number of successes divided by test trials) in ascending order, where the largest proportion was ranked 80, reflecting thus the highest performance in our fish (sample size is 80 fish).

#### 2- Associative and reversal learning tasks

In the associative learning task, we exposed the fish to a two-colour choice test to estimate fish learning abilities through operant conditioning in associating a food reward with a colour cue, yellow vs red. In the test compartment, we placed two 1 mL plastic pipettes covered with either yellow or red adhesive tape, and each contained a defrosted adult artemia. An opaque grey Plexiglas rectangle plate separated the two pipettes creating thus two zones of choice. A test trial started with the experimenter pulling up the opaque sliding door followed by the see-through door, giving the fish a few seconds to see the set-up before entering the test compartment and choosing one of the two pipettes. The experimenter scored a choice as “correct” if a fish entered with its body length the zone of the rewarding colour at its first attempt and “failure” if it chose the non-rewarding colour at its first attempt (see Supplementary Videos). We balanced the colour and side of the rewarding pipette across fish and trials. As such, half of the fish had red as the rewarding colour while the other half had yellow with 50-50 presentation of the rewarding colour on the left and right side of the test compartments (with no more than three presentations on the same side in succession) (following on the protocol by Triki et al. (2022b) in guppy fish).

Once a fish learned the colour-reward association in the associative learning phase, we tested its abilities in a reversal learning phase. It consisted of reversing the reward contingency, and the previously unrewarding colour became the new rewarding cue. For example, if a fish learned the yellow-reward association in the previous task, in the reversal task, it had to learn the red-reward association instead. For associative and reversal learning phases, the fish received a total of twenty test sessions with six trials daily. We set the learning criteria in each test to a score of either six correct choices out of six consecutive trials or five correct choices out of six trials in two consecutive sessions. The probability of learning by chance with these criteria is p < 0.05 (with a binomial test).

Finally, for the associative learning performance, we ranked fish success and the number of sessions needed to learn the task in descending order, where the smallest number of sessions to succeed was ranked 80, reflecting thus the highest performance in our fish. In the reversal learning performance, we ranked fish success and the number of sessions needed to pass first the associative learning phase and then the reversal phase in descending order, where the smallest number of sessions to succeed was ranked 80, reflecting thus the highest performance in our fish (sample size is 80 fish).

### Brain staining and 3D-image acquisition

We prepared the 29 female guppies from the baseline group and 80 from the treatment group for X-ray brain scans by first euthanizing them with an overdose of benzocaine (0.4 g L^-1^). With a digital calliper, we estimated the fish body size as standard length (SL) to the nearest 0.01 millimetre. We then fixated their whole heads in 4% paraformaldehyde phosphate-buffered saline (PBS) for five days. After that, we washed the samples twice in PBS for 10 min and kept the samples in PBS. We then followed the Phosphotungstic acid (PTA) staining protocol by Lesciotto et al. (2020) to prepare our samples for X-ray scanning. In this protocol, we first dehydrated the samples by placing them in a series of solutions as follows: one day in 30% ethanol in PBS; one day in 50% ethanol in PBS; one day in 70% ethanol in PBS; one hour in a solution with a ratio of 4:4:3 volumes of ethanol, methanol, and H2O; one hour in 80% methanol in H2O; and one hour in 90% methanol in H2O. After that, we proceeded with the staining phase by placing the samples in 0.7% PTA in 90% methanol in H2O for 23 days. Twenty-three days was the optimal staining duration for our samples based on pilot rapid X-ray scans to check the staining quality (see below). We then rehydrated the samples by placing them in 90% methanol for six hours; 80% methanol overnight; 70% methanol for six hours; 50% methanol overnight; 30% methanol for a day; in PBS for one day; and finally storage in 0.01% sodium azide in PBS.

We transferred the samples to the Stockholm University Brain Imagery Center (SUBIC) for image acquisition. We scanned the samples using a Zeiss Xradia Versa 520, with the X-ray source at a voltage of 100 kV and a power of 9W. We used the 0.4x objective coupled with a scintillator. The source-to-sample distance was 30 mm and sample-to-detector distance was 81 mm. The effective voxel size was 9.17 μm with a compensated optical and geometrical magnification. The scan consisted of 1201 projections over 360 degrees with 1s exposure time for each projection. Each scan took 1 h and 36 min on average including reference images and readout time of the CCD camera. Given the small size of fish heads and to optimise the scan time, we arranged four samples per scan (see Supplementary Material). We obtained three-dimensional images of the brain scans through an automatised tomography reconstruction with Zeiss Scout-and-Scan software.

### Brain morphology measurements

To segment the 3D brain images, we first aligned the images digitally in Dragonfly (Dragonfly 2020.2 [Computer software] 2020) in three planes – transversal, coronal, and sagittal to the cardinal axes (Fig. 1). To ensure accuracy, we either adjusted the voxel size of the dataset through resampling using bicubic interpolation or made note of any changes in voxel size and corrected the volume accordingly. This was necessary due to the potential for minor changes in voxel sizes caused by alignment. We obtained full head images in the scans and then cropped them to only include the brain tissue. This was done consistently across all planes to improve segmentation and accurately measure the volumes.

For the segmentation *per* se, we first generated a semi-manually segmented brain into five regions (telencephalon, mesencephalon, diencephalon, cerebellum and brain stem) (Fig. 1). This was achieved by employing Biomedisa (Lösel et al. 2020) using random walker interpolation between sparsely manually segmented slices. In total, we semi-manually segmented 23 samples and used them as an Elastix template (Klein et al. 2009). We then manually checked these samples and corrected potential errors, then used them as a training dataset for the following deep-learning-based segmentation on the rest of the samples. We used the deep-learning algorithm from Schoppe et al. (2020), which is U-net-like (Ronneberger et al. 2015). Finally, we computed the brain regions’ volumes by multiplying the voxel number and voxel size from the segmented labels.

### Data analysis

We used the open-access software R, version 4.2.1 (R Core Team 2022), to run all statistical analyses and generate the figures. Overall, we ran four different statistical analysis approaches. First, we tested whether the fish exposed to different social treatments may have developed different cognitive abilities. We analysed the rank performance data for all three abilities: inhibitory control, associative learning, and reversal learning. Given that this data violates the overdispersion assumption for count data, we fitted three generalised linear mixed models using a template model builder (glmmTMB) with a negative binomial distribution. In one model, we fitted inhibitory control rank performance as the dependent variable, social treatment as the predictor and batch as a random factor. For the associative and reversal learning models, we fitted rank performance as the dependent variable, treatment as a predictor, and the colour of the pipette as a covariate to control statistically for potential colour bias, while the batch was the random factor.

Second, we tested whether brain morphology (total brain size and brain region sizes) differed across the three social treatments in both the baseline and test groups. To do so, we ran two linear mixed effect models (LMMs), one for baseline data and one for test data, where we fitted log-transformed brain size (mm^3^) as the dependent variable, social treatment (with three levels: three, three guppies plus three splash tetras, and six) as the fixed predictor, log-transformed and standardised body size (mm) as a covariate, and batch number (we had two batches, see above) as a random factor. For the five brain regions, telencephalon, diencephalon, mesencephalon, cerebellum and brain stem, similarly, we fitted two multivariate analyses of variance (MANOVA) for baseline and test data. We fitted the dependent variable as a matrix of all five brain region sizes log-transformed and standardised (e.g., (Triki et al. 2019)), with treatment as a predictor and body size as a covariate. We also fitted batch as a predictor since MANOVA does not support mixed effects.

Third, we tested whether cognitive performance was influenced by individual brain size and region size. In preparation for these analyses, we extracted the residuals of log brain size on log body size, as well as the residuals of every brain region size (log-transformed) on log-transformed and standardised brain and body sizes. Then, we fitted a set of glmmTMB models with the designated cognitive ability as the dependent variable while social treatment and the designated brain measurement as predictors. We also fitted batch as the random factor in all these models, while for the models testing for associative and reversal learning, we added the colour of the pipette as a co-variate.

Finally, we checked that the fitted models met their corresponding assumptions, such as normality of residuals and homogeneity of variance. For further details, we provide a step-by-step code and data used to generate the findings in the present study (see the Data and Code accessibility statement).

## Ethics

This work was approved by the ethics research committee of the Stockholm Animal Research Ethical Permit Board [permit numbers: Dnr 17362-2019, 17402-2020].

## Data availability

Source data from this study are archived in the Figshare data repository DOI: 10.6084/m9.figshare.23500839.

## Code availability

Analyses reported in this article can be reproduced using code archived at the Figshare data repository DOI: 10.6084/m9.figshare.23500839.

## Author contribution

ZT conceived the idea. ZT and NK designed the study. ZT and EA collected the data. ESN and OS analysed the video recordings. TZ segmented the 3D brain scans with input from ZT. ZT analysed the data, generated the figures, and wrote the manuscript with input from NK and TZ. All authors gave final approval for publication and agreed to be held accountable for the work performed therein.

## Acknowledgement

We thank Mirjam Amcoff for all the help and support with the logistics needed for this study. This work was supported by the Swiss National Science Foundation [grant numbers P2NEP3_188240, P400PB_199286 and PZ00P3_209020 to ZT] and the Swedish Research Council [grant numbers 2016-03435 and 2021-04476 to NK]. Brain data acquisition was supported by a grant from the Stockholm University Brain Imaging Centre [grant number SU FV-5.1.2-1035-15 to ZT].

